# A dual prokaryotic (*E. coli*) expression system (pdMAX)

**DOI:** 10.1101/2020.07.23.217315

**Authors:** Manabu Murakami, Agnieszka M. Murakami, Kazuyoshi Hirota, Shirou Itagaki

## Abstract

In this study, we introduced an efficient subcloning and expression system with two inducible prokaryotic expression promoters, arabinose and lac, in a single plasmid in *Escherichia coli*. This pdMAX system is a manageable size at 5811 bp. The arabinose promoter unit allows for the expression of a FLAG-tagged protein, while the isopropyl-β-D-thiogalactoside (IPTG)-inducible unit allows for the expression of a Myc-tagged protein. An efficient subcloning (DNA insertion) system (iUnit) follows each promoter. The iUnit, based on a toxin that targets DNA topoisomerase of *E. coli*, allows for effective selection with arabinose or IPTG induction.

Interestingly, the dual induction plasmid system shows limited protein expression. To analyze the expression of inserted genes, we performed an *α*-complementation assay, in which the *α*-peptide of the lacZ (β-galactosidase) gene is inserted in XL-10 *E. coli* cells. *E. coli* expressing the recombinant plasmid form blue colonies when plated on ampicillin/IPTG- or arabinose/X-gal-containing plates if the *α*-peptide was correctly inserted in-frame to produce the tagged protein. This assay system assesses whether the *α*-peptide was inserted at the restriction sites (EcoRV or SmaI), whether the inserted peptide was expressed, and whether the inserted *α*-peptide sequence was ligated in-frame to produce a tagged protein. Frameshifts result in no *α*-complementation and white colonies.

With the dual promoter plasmid (pdMAX) system, expressed *lacZ* activity was significantly decreased comparing with the solo expression (pgMAX) system. Despite this disadvantage, we believe the pdMAX system is still useful for the analysis of distinct genes in *E. coli*, which will enable different types of expression analysis. Overall, the novel pdMAX system allows for efficient subcloning of two different genes. Furthermore, the pdMAX system could be used to induce and analyze the expression of two distinct genes and adopted to various types of prokaryotic gene expression analyses.

## Introduction

A number of commercial bacterial expression plasmids are used for prokaryotic expression in *Escherichia coli*. At present, the use of different but compatible vectors is commonly used for the simultaneous expression of two distinct proteins [1]. For example, the pET vector can be used with the pRSET vector (Novagen, Merck KGaA, Darmstadt, Germany), which contain different origins of replication (pBR322 and pUC origins, respectively). Because of different plasmid copy numbers (the pRSET vector is high copy number but pET is not), it is possible to obtain significantly more protein expression from pRSET than from pET in *E. coli*. As two-plasmid systems have the disadvantage of unstable plasmid copy numbers, a single plasmid system is needed.

We previously established a novel solo expression (pgMAX) system, a dual expression system with two (prokaryotic and mammalian) expression modes [2]. This novel pgMAX system enabled efficient subcloning and gene expression in *E. coli.* Furthermore, this system enabled rapid construction of a mammalian expression plasmid with a simple deletion step and is referred to as a deletion (D)-type system.

In the present study, we established a dual promoter expression plasmid for use in *E. coli*. This novel dual expression (pdMAX) system has two inducible promoters, arabinose and isopropyl-β-D-thiogalactoside (IPTG). To analyze the expression of inserted genes, we examined insertion of the *α*-peptide sequence of the *lacZ* gene, commonly referred to as the *α-*complementation assay, which enabled detection of *lacZ* activity via the formation of blue colonies [3]. The novel pdMAX system simplifies inducible prokaryotic gene expression analyses and will be useful for expression analysis of different proteins in *E. coli*.

## Methods

### Plasmid construction

The novel pdMAXflag/myc originated from pgMAX [2]. The pBluescript plasmid was purchased from Agilent Technologies (Santa Clara, CA, USA). The pdMAX plasmid was constructed by serial PCR amplification and ligation. The araC gene and araBAD promoter were PCR-amplified from pBAD/His/lacZ (Invitrogen, Carlsbad, CA, USA). The lac promoter and inhibitory unit (iUnit) were PCR-amplified from pgMAX. The final plasmid sequence is shown in Supporting Information Figure 1. Primers used in the construction of pdMAX are provided in Supporting Information Table 1. PCR conditions using high-fidelity Pfu DNA polymerase (Agilent Technologies) were as follows: 35 cycles of denaturation at 94°C for 20 s, annealing at the calculated temperature for 30 s, and extension at 72°C for 30 s. Amplified PCR products were gel-purified with a gel extraction kit (Macherey-Nagel GmbH, Dueren, Germany). Protocols using the pdMAX system (dx.doi.org/10.17504/protocols.io.binfkdbn) were based upon the former pgMAX system, which have been deposited in protocols.io (DOI dx.doi.org/10.17504/protocols.io.zq3f5yn).

### α-Complementation assay

The host *E. coli* strain (XL-10) carries the *lacZ* deletion mutant (*lacZΔM15*), which contains the ω-peptide [3]. If the recombinant plasmid carries the *lacZ α-*sequence, which encodes the first 59 amino acids of β-galactosidase (the α-peptide), the ω- and α-peptides will be expressed together, resulting in a functional β-galactosidase enzyme [4, 5]. Based on this, we inserted a PCR-amplified α-peptide sequence (LQRRDWENPGVTQLNRLAAHPPFASWRNSEE) into blunt-end restriction enzyme sites (EcoRV or SmaI). If the α-peptide sequence is inserted in frame with the FLAG (arabinose unit) or myc (IPTG unit) tag sequences, and in the sense direction, the α-peptide is expressed following induction with arabinose or IPTG, resulting in the formation of blue colonies on X-gal-containing lysogeny broth (LB) plates.

For the *α*-complementation assay, a blunt-end DNA fragment (369 bp) of the *α*-peptide sequence of the *lacZ* gene was amplified using Pfu DNA polymerase with specific primers (AlphaFor, CAGGAAACAGCTATGAC; AlphaRev, CCATTCGCCATTCAGGCTGCGCAA) and pBluescript KS^-^ plasmid (Agilent Technologies) as a template. The PCR-amplified product was inserted at the EcoRV (arabinose promoter) or SmaI (IPTG promoter) site of pdMAX.

DNA ligation was performed using standard ligation techniques (Takara DNA Ligation kit ver. 2.1; Takara, Otsu, Japan). For transformation, XL10-Gold ultracompetent cells (Tet^r^ Δ(mcrA)183 Δ(mcrCB-hsdSMR-mrr)173 endA1 supE44 thi-1 recA1 gyrA96 relA1 lac The [F’proAB lacI^q^ZΔM15 Tn10 (Tet^r^) Amy Cam^r^]; Agilent Technologies) were used. After 16 h of incubation on LB agar plates containing ampicillin (ABPC; 150 μg/mL), X-gal (1 mM), and IPTG (1 mM) or arabinose (10 mM), colonies were observed.

### Spot culture of recombinant clones

Confirmed by restriction analysis after small-prep DNA analysis, recombinant clones were diluted 100 times with LB medium and 2 μL of each clone was spotted on ABPC (with/without arabinose, IPTG, with/without X-gal)-containing plates. After 16 h at 37°C, colonies were observed.

### Evaluation of blue colonies

To measure enzymatic activity of *lacZ, E. coli* cells with the recombinant plasmid were incubated in LB medium (0.5 mL) with X-gal (1.0 mM) on an orbital rocker at 37°C for 12 h with varying concentrations of IPTG (0.01–1 mM) or arabinose (0.3–30 mM). Accumulation of blue-colored precipitates, indicating β-galactosidase activity, was evaluated at 490 nm (GENESYS 10S UV-Vis; Thermo Fisher Scientific, Waltham, MA, USA).

### Statistical analysis

Data are expressed as the means ± standard error of the mean. Prior to statistical analyses, data were analyzed with the Shapiro–Wilk test. After confirmation of a normal distribution, statistical differences were further determined by Student’s *t*-test. *P* < 0.05 was considered to indicate statistical significance.

## Results

### Plasmid construction

Figure 1A shows the plasmid map of pdMAX, which was based on pgMAX [2]. The novel pdMAX plasmid includes arabinose- and IPTG-inducible promoters. The arabinose-inducible promoter is composed of araC and the BAD promoter, which originated from the commercially available pBAD/His plasmid (Invitrogen). The IPTG-sensitive promoter is composed of the lac promoter and lac operator. Both promoters were followed by the inhibitory unit iUnit for efficient subcloning, as previously described [2]. The arabinose unit was composed of araC, araBAD, the FLAG tag sequence, EcoRV site for blunt-end cloning, and iUnit (Figure 1A) while the IPTG unit was composed of the lac promoter, lac operator, the myc tag sequence, SmaI site for blunt-end cloning, and iUnit. The iUnit, which originates from CcdB, a toxin targeting the essential DNA gyrase of *E. coli* [6], enables efficient subcloning, as plasmids with no insert will not form colonies.

**Figure 1.**
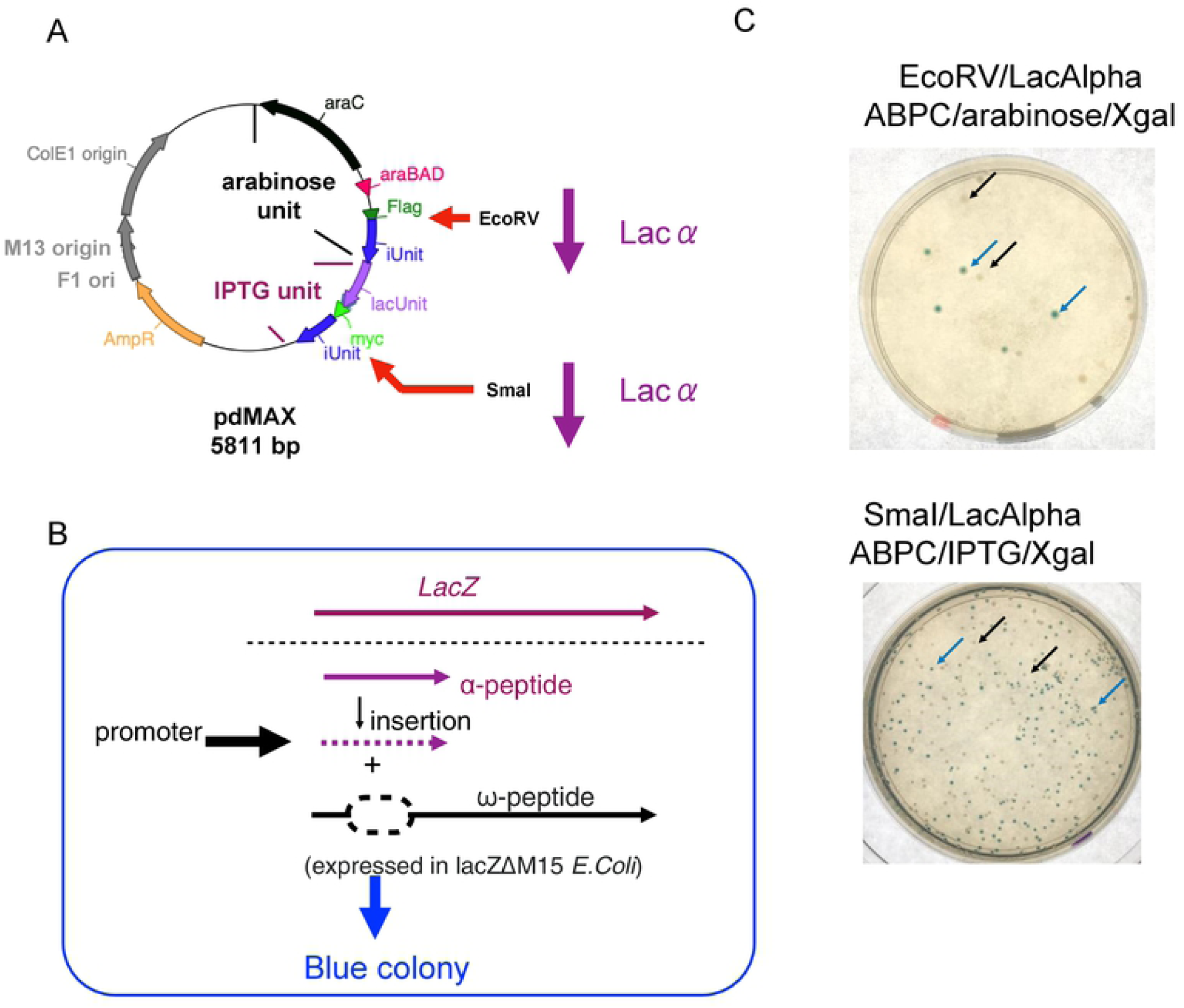
The pdMAX plasmid system. A. pgMAXs/FLAG construct. The pdMAX system has two functional expression units, arabinose and IPTG. The arabinose unit contains araC (black arrow), araBAD promoter (red triangle), Kozak sequence, FLAG-tag sequence (green triangle), EcoRV site, and iUnit (blue arrow). The IPTG unit contains the lac unit (lac promoter and lac operator, lavender arrow), Kozak sequence, myc-tag sequence (light green triangle), SmaI site, and iUnit (blue arrow). The restriction enzyme and subcloning (EcoRV and SmaI) sites for the α-peptide sequence (lac-α, purple arrow) are indicated. B. Scheme of the α-complementation assay. PCR-amplified blunt-ended α-peptide sequence was inserted into the cloning site under the control of the inducible promoter. Following induction, the α-peptide is expressed along with the ω-peptide, resulting in α-complementation of *lacZ* (β-galactosidase). With active β-galactosidase, X-gal (a colorless analog of lactose) is cleaved to form an insoluble blue pigment. C. α-Complementation assay. PCR-amplified α-peptide was inserted into the EcoRV (arabinose unit, upper panel) or SmaI (IPTG unit, lower panel) sites. Recombinant clones with lacZ α-peptide insertion in the sense direction resulted in blue colonies (blue arrow), while clones without the insert or insertion in the antisense direction resulted in white colonies (black arrow). Approximately 30–40% of the clones formed blue colonies, indicating a high insertion rate with the pdMAX system.

To confirm protein expression, the *α*-peptide sequence of the *lacZ* gene was PCR amplified, purified, and subcloned into blunt-end sites of EcoRV (arabinose unit) or SmaI (IPTG unit) in pdMAX. Following ligation and transformation, the recombinant clones were plated on LB agar containing ABPC (150 μg/mL), X-gal (1 mM), and arabinose (10 mM, for arabinose unit induction) or IPTG (1 mM, for lac operon induction). After 16 h, colonies were observed and numbers of blue (*α*-complementation) or white (antisense directed ligation or no insertion of the DNA fragment) colonies were evaluated.

Figure 1C shows representative images of α-complementation at EcoRV (upper panel) and SmaI (lower panel) sites. Approximately 30–40% of the colonies were blue, suggesting efficient subcloning of the *α*-peptide sequence, whereas no dephosphorylation with alkaline phosphatase was applied for the vector (pdMAXs), which is a common step in traditional cloning workflows to ensure that the vector does not re-circularize during ligation. It is expected that approximately 50% of the fragments will insert in the sense direction. Therefore, ligation efficiency should be high at both the EcoRV (arabinose-sensitive unit) and SmaI (IPTG-sensitive unit) sites.

### Spot culture of recombinant clones

*α*-Complementation can result in white or light blue colonies. Therefore, a secondary blue–white screening of the target colonies is required [5]. To confirm expression of the *α*-peptide or iUnits, inserted after each cloning site (EcoRV or SmaI), we examined secondary spot cultures of recombinant clones, which are modified *E. coli* cultures as described by Zhang [5]. If *α*-complementation occurs on X-gal-containing plates, blue colonies are formed. If the iUnit is expressed, no colonies are formed [2]. Figure 2 shows typical images of different culture conditions. On ABPC-containing plates, pBluescript-containing clones showed white colonies (Figure 2, lane 1, ABPC plate). Other clones of pgMAXs (IPTG-inducible iUnit-containing plasmid, lane 2), pgMAXs with *α*-peptide (lane 3), pdMAX (lane 4), pdMAX with *α*-peptide in EcoRV (lane 5), and pdMAX with *α*-peptide in SmaI (lane 6) also formed white colonies, while XL-10 cells did not form any colonies (negative control, lane 7). Only pBluescript and pgMAXs with *α*-peptide formed blue colonies on ABPC- and X-gal-containing plates (ABPC/X-gal lanes 1 and3), while pdMAX with the *α*-peptide at EcoRV or SmaI sites formed white colonies, indicating marginal expression of the *α*-peptide in both clones without gene induction (ABPC/X-gal, lanes 5 and 6). Arabinose induction resulted in blue colony formation in pBluescript and pdMAX with the *α*-peptide at the EcoRV site (ABPC/X-gal/arabinose, lanes 1 and 5), indicating induction of the *α*-peptide with the arabinose unit. Meanwhile, arabinose induction resulted in no colony formation with pdMAX and pdMAX with the *α*-peptide inserted at the SmaI site, indicating expression of iUnit in these transformants (lanes 4 and 6). IPTG induction showed blue colonies in pBluescript, pgMAXs with the *α*-peptide, and pdMAX with the *α*-peptide at SmaI (ABPC/X-gal/IPTG, lanes 1, 3, and 6), indicating expression of the *α*-peptide and *α*-complementation. IPTG induction resulted in no colony formation in pgMAX, pdMAX, and pdMAX with the *α*-peptide at the EcoRV site (lanes 2, 4, and 5), indicating expression of iUnit. Taken together, spot culture analysis showed induction of arabinose and IPTG units. Interestingly, only pBluescript and pgMAX with the *α*-peptide showed blue colonies on X-gal-containing plates, while pdMAX with the *α*-peptide at EcoRV or SmaI resulted in white colonies, indicating low expression of the *α*-peptide in these two clones. Considering the sensitivity of the *lacZ* system, our data suggest that the pdMAX system results in low basal expression of the arabinose and IPTG units.

**Figure 2.**
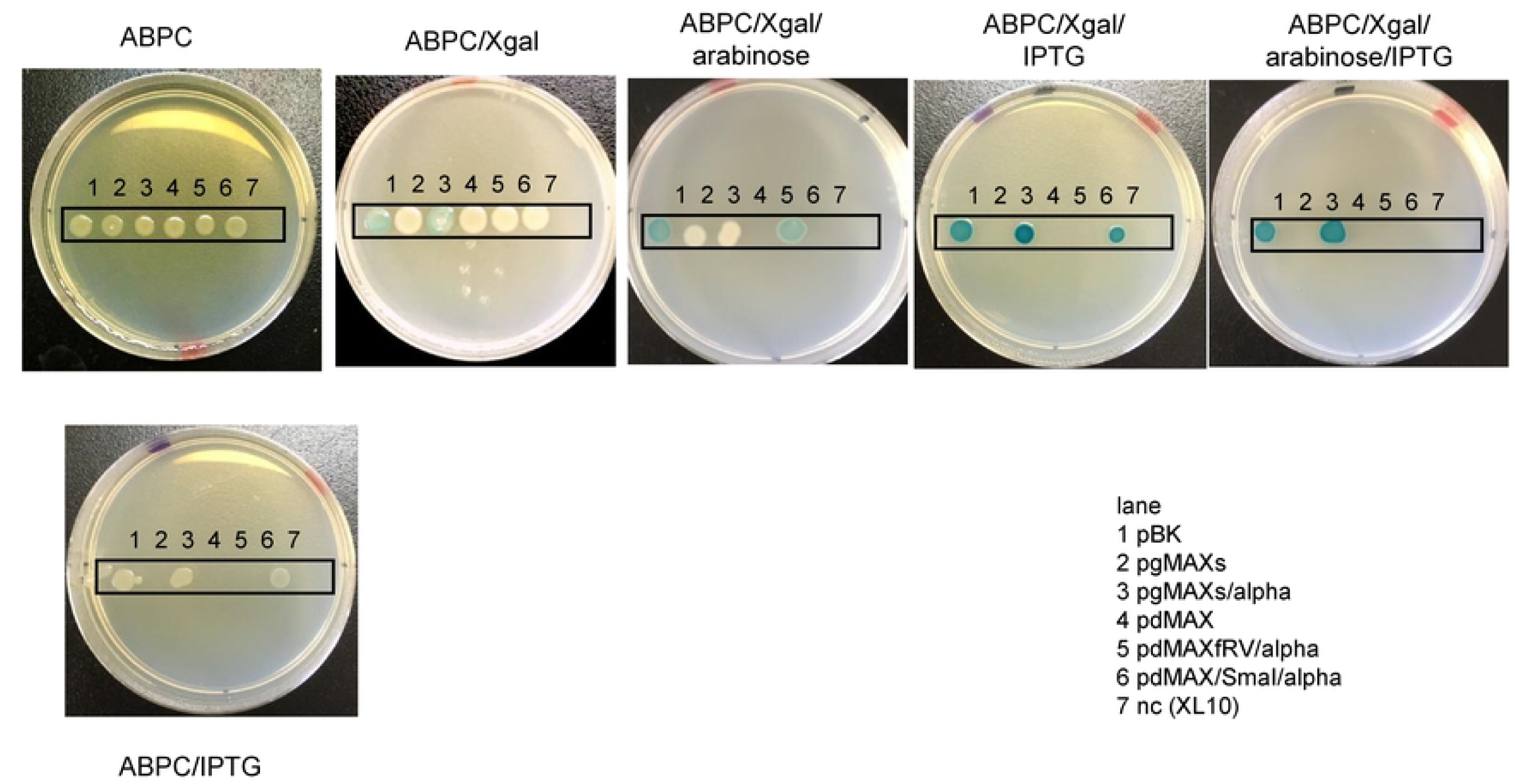
Spot culture and α-peptide induction. Various types of plasmids were diluted (x100) and 2 μL of culture was spotted on LB plates in various conditions. Lane 1, pBluescript (pBK); lane 2, pgMAXs; lane 3, pgMAXs/α-peptide; lane 4, pdMAX; lane 5, pdMAX with α-peptide at the EcoRV site (arabinose unit); lane 6, pdMAX with α-peptide at the SmaI site (IPTG unit); and lane 7, negative control (nc; XL-10 cells). Conditions of the LB plates were as follows: ABPC (50 μg/mL), X-gal (1 mM), arabinose (10 mM), and IPTG (1 mM). In ABPC/X-gal-containing plates, only pBluescript and pgMAXs/α-peptide formed blue colonies, while pdMAX with the α-peptide formed white colonies (lanes 5 and 6). In ABPC/X-gal/arabinose-containing plates, pdMAX with the α-peptide at the EcoRV site formed blue colonies (lane 5), while pdMAX and pdMAX with the α-peptide at the SmaI site did not form any colonies (lanes 4 and 6, respectively). In ABPC/X-gal/IPTG-containing plates, pdMAX with the α-peptide at the SmaI site did not form colonies (lane 6).

To analyze protein expression in the pdMAX system, genes labeled with enhanced green fluorescent protein (EGFP) or DsRed were inserted in the arabinose (EcoRV site) or IPTG (SmaI site) units. Insertion of DsRed or EGFP in the IPTG unit resulted in no apparent fluorescent protein expression (Supporting Information Figure 2), although ligation at the SmaI site showed efficient subcloning (about 80%), indicating low protein expression in the pdMAX plasmid system. Insertion in the arabinose unit at the EcoRV site was also successful (about 80%). Successful subcloning at both sites indicated some expression of iUnit in both cases; otherwise, the subcloning rate of blunt-end DNA fragments would be low. We also examined DsRed fluorescence after successful subcloning at the EcoRV site (arabinose unit); no fluorescence was observed (data not shown). However, insertion of EGFP or DsRed in the pgMAX plasmid, which has only a single IPTG unit, resulted in high expression of fluorescent proteins (Supporting Information Figure 2), as previously reported [2].

### Dose-dependent induction of lacZ-α-peptide expression with arabinose or IPTG

Figure 3A shows concentration-dependent color changes due to pdMAX/EcoRV/A with the *α*-sequence inserted in the arabinose unit and pdMAX/SmaI/A with the *α*-sequence inserted in the IPTG unit. The pBluescript plasmid showed significant changes in blue color in response to IPTG (pBK). Arabinose resulted in limited expression of X-gal-related blue color, while IPTG induced significant color changes (pdMAX/SmaI/A). Statistical analysis showed concentration-dependent changes in blue color with the three constructs (Figure 3B). Taken together, the pdMAX plasmid resulted in efficient subcloning. In addition, dual inducible promoters (arabinose and IPTG) enabled concentration-dependent *lacZ*-*α*-peptide expression.

**Figure 3.**
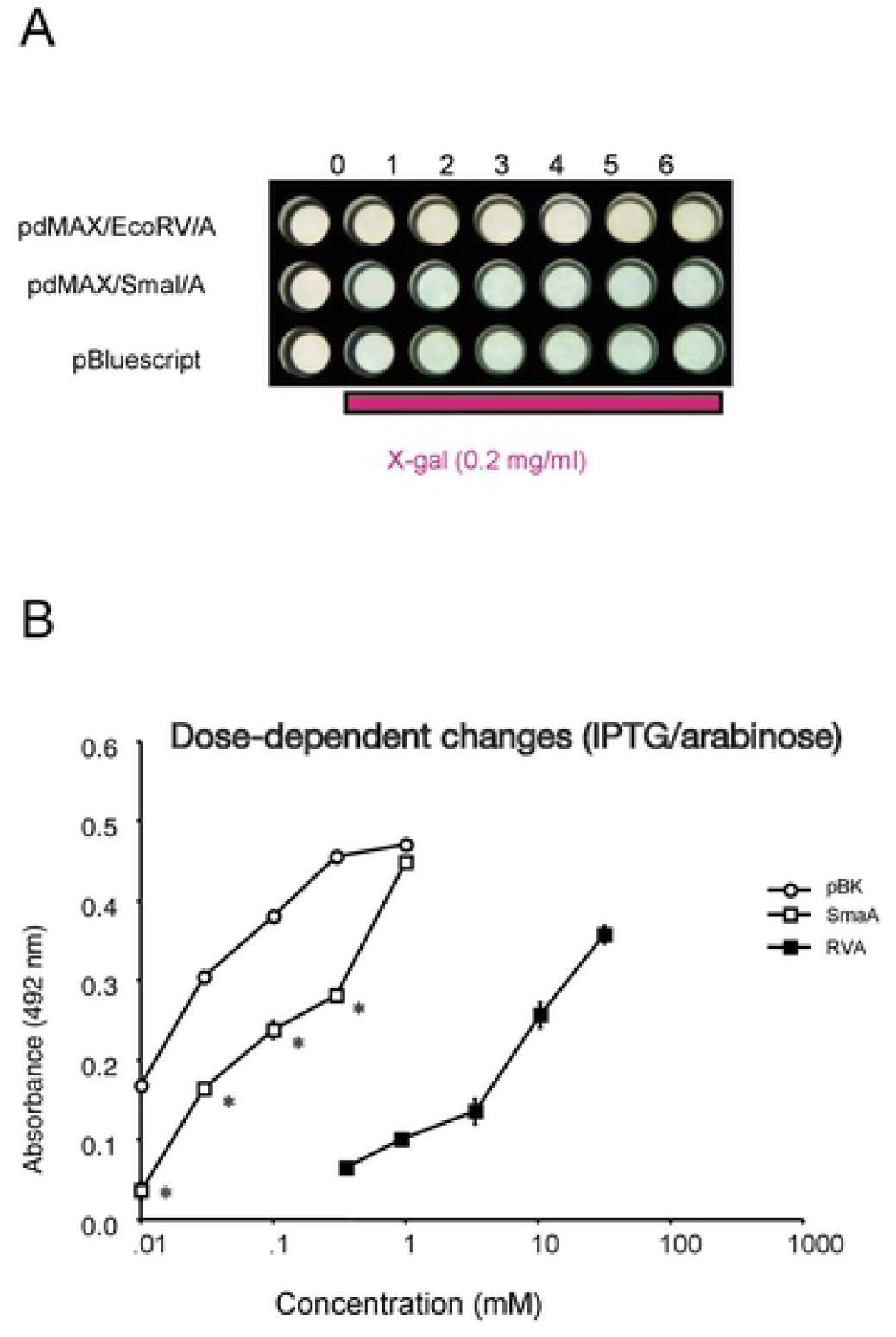
Dose-dependent induction of α-peptide induction. A. Representative image of *lacZ*-related blue *E. coli* clones. Clones containing the recombinant plasmid with the α-peptide were inoculated in 0.5 mL of LB medium with ampicillin, X-gal (1 mM), and various concentrations of IPTG or arabinose. Lane 0, no X-gal, IPTG, or arabinose; lane 1, X-gal; lane 2, 0.01 mM IPTG or 0.3 mM arabinose; lane 3, 0.03 mM IPTG or 1 mM arabinose; lane 4, 0.1 mM IPTG or 3 mM arabinose; lane 5, 0.3 mM IPTG or 10 mM arabinose; and lane 6, 1 mM IPTG or 30 mM arabinose. B. Statistical analysis of *lacZ*-related blue color. Accumulated blue color was measured at 490 nm. *E. coli* with pBK showed IPTG dose-dependent blue colonies (open circle). Recombinant clones of pdMAX/SmaI/α-peptide (SmaA) also showed dose-dependent blue color following induction with IPTG, although the blue color was significantly reduced (open square). Recombinant clones of pdMAX/EcoRV/α-peptide (RVA) also showed arabinose concentration-dependent blue color (closed square). N = 6. *p < 0.05 vs. pBK.

## Discussion

In the present study, we established a dual inducible subcloning/expression plasmid (pdMAX). This plasmid enables simple and efficient subcloning of desired genes with standard techniques, and regulated induction of genes with arabinose or IPTG. EcoRV and SmaI restriction sites in the arabinose and IPTG unit, respectively, are useful for blunt-end DNA fragments, such as PCR-amplified products generated with robust high-fidelity DNA polymerases, which have exonuclease activity for accurate PCR amplification; we used Pfu in this study.

Although we achieved efficient subcloning in the arabinose and IPTG units in pdMAX, expression levels of the inserted genes were low compared with previous reports using the single expression construct pgMAX [2]. Despite the low protein expression with the arabinose and IPTG units, we would like to emphasize the possible applications of the independent induction of the expression of two distinct proteins, such as the study of protein–protein interactions and the inhibitory effects of one protein on another.

In the present study, we applied two different induction systems, arabinose and IPTG units. The arabinose unit is composed of AraC and the araBAD promoter (pBAD), which is arabinose sensitive; AraC is a regulatory protein for pBAD. In the presence of arabinose, AraC positively regulates pBAD promoter activity, whereas in the absence of arabinose, AraC tightly represses target gene expression [7]. The regulatory elements of the *E. coli* arabinose operon in the pdMAX system made it possible to precisely modulate gene expression levels, allowing optimization of the yields of the protein of interest.

The IPTG unit is composed of the lac promoter (lacP) and operator (lacO). IPTG is frequently used as an inducer of the *lac* operon [8]. As IPTG cannot be metabolized by *E. coli*, the concentration of IPTG remains constant, resulting in constant expression of the *lac* promoter and operator-controlled genes. Thus, the inserted *α*-peptide of lacZ (β-galactosidase) was thought to be constitutively expressed with IPTG induction. While IPTG-sensitive systems are known to exhibit some basal expression, pdMAX has relatively low expression in the basal state (Figure 2, ABPC/X-gal, and Figure 3B), indicating titratable expression of the inserted gene product.

To express two proteins in *E. coli* simultaneously, different but compatible vectors are commonly used; however, regulation of expression is difficult. To regulate the expression of two independent proteins, the pdMAX plasmid might be advantageous. With the regulation afforded by this novel system, it might be possible to analyze the relationship between different proteins, i.e., interactions or inhibitory effects of different proteins.

As a single-plasmid strategy, the pET Duet plasmid is commonly used. This system consists of the T7 promoter expression vector, designed to co-express two target proteins in *E. coli* by IPTG induction. Therefore, both proteins cannot be induced independently [9]. In addition, pET Duet requires a bacterial strain that expresses T7 RNA polymerase, such as the BL21 (DE3) strain. If we consider to make expression analysis with cDNA-library, XL-10 cells have several advantages, such as high efficiency for large plasmids, the ability to clone methylated DNA, and high-quality mini-prep plasmids.

In the present study, we performed an α-complementation assay to assess induction of protein expression on X-gal-containing LB plates. While this assay is simple, there are several disadvantages. The α-complementation assay is influenced by various factors, such as PCR conditions, ligase quality, competent cell quality, quality of vector preparation, and molar ratio between the insert and vector, as it depends on DNA ligation of the PCR product. Furthermore, direction of blunt-end DNA fragment ligation cannot be controlled; thus, it cannot be expected to yield 100% blue colony formation. Following α-peptide ligation and transformation, four types of clones are expected: self-ligation of the vector plasmid, insertion of incorrectly amplified PCR fragment, insertion of correctly amplified PCR fragment in the antisense direction, and insertion of correctly amplified PCR fragment in the sense direction. Only the final case results in the formation of blue colonies. As a result, only 30–40% of recombinant clones with the original pgMAXs formed blue colonies (Figure 1). We also have to consider possible white (or light blue) colony formation with correct (sense-directed) insertion of the α-peptide sequence. In spite of these disadvantages, the α-complementation assay was effective, as several hundred colonies could be easily analyzed.

In the present study, we used an iUnit from CcdB, a toxin targeting the essential DNA gyrase of *E. coli* [6]. As iUnit selection is effective for fragment ligation (approximately 80% of colonies have the insert), pdMAX has significant superiority over classical DNA expression systems. However, we believe it is still possible to improve the iUnit, e.g., with the use of short-peptide toxins. If a short-peptide toxin could be used, it might be possible to form a chimeric construct of two proteins.

We focused on a prokaryotic expression plasmid for *E. coli*, which is commonly used to replicate plasmids. *E. coli* is also commonly used for protein production, as well as analyses of protein–protein interactions and DNA–protein interactions, and functional analyses of cloned genes. Therefore, expression of two proteins with different induction systems using pdMAX could be applied for various types of analysis.

## Conclusion

We established a subcloning and expression plasmid system for two different genes in *E. coli*. The pdMAX plasmid system enables highly efficient subcloning of blunt-end DNA fragments and inducible expression with arabinose or IPTG.

## Acknowledgments

This research was sponsored in part by Grants-in-Aid for Scientific Research from the Japan Society for the Promotion of Science (KAKENHI nos. 17K08527 [MM], 17H04319 [KH], and 20K07255 [MM]). No additional external funding was received for this study. We thank Mr. Maximilian Murakami for his technical advice.

## Author contributions

Experiments were conceived and designed by MM. Experiments were performed by MM, AMM, KH, and SI. Data analyses were performed by AMM and MM. Reagents, materials, and analysis tools were provided by SI and KH. The manuscript was written by MM, AMM, KH, and SI.

## Additional information

We declare no competing financial interests.

## Figure legends

**Supporting Information Table 1. Primers used in the construction of pdMAX.**

**Supporting Information Figure 1. Sequence of pdMAX.**

**Supporting Information Figure 2. Insertion of fluorescent protein sequences.**

A. Fluorescence of pgMAX/DsRed (R), pgMAX/EGFP (E), pdMAX/IPTG/EGFP (1), pdMAX/IPTG/DsRed (2), and pgMAX (N; negative control). Colonies of pgMAX/DsRed (R), pgMAX/EGFP (E), pdMAX/IPTG/EGFP (1), pdMAX/IPTG/DsRed (2), and pgMAX (N) were observed under a white light (upper panel). Fluorescence was observed under a blue light and orange filter (excitation wavelength, 470 nm; emission wavelength, 505 nm).

B. Expression of fluorescent proteins with the pgMAX system. Recombinant clones of the pgMAX/blunt-end DsRed fragment (R in red), pgMAX/blunt-end EGFP fragment (E in green), negative control (pdMAX), pdMAX/blunt-end EGFP fragment in IPTG unit (1 in green), and pdMAX/blunt-end DsRed fragment (2 in red) are shown. Recombinant clones were transformed into *E. coli* and incubated in LB medium supplemented with ABPC and IPTG (pgMAX/DsRed, pgMAX/EGFP, pdMAX/IPTG/EGFP, and pdMAX/IPTG/DsRed). Transformed *E. coli* were collected by centrifugation (5000 × *g*, 1 min) and observed under a white light (upper panel), or blue light and orange filter (lower panel, excitation wavelength, 470 nm; emission wavelength, 505 nm). Note that the pdMAX plasmid showed marginal fluorescence with EGFP (1) and DsRed (2).

